# Serotonergic and dopaminergic neurons in the dorsal raphe are differentially altered in a mouse model for parkinsonism

**DOI:** 10.1101/2023.07.21.550014

**Authors:** Laura Boi, Yvonne Johansson, Raffaella Tonini, Rosario Moratalla, Gilberto Fisone, Gilad Silberberg

## Abstract

Parkinson’s disease (PD) is characterized by motor impairments caused by degeneration of dopamine neurons in the substantia nigra pars compacta. In addition to these symptoms, PD patients often suffer from non-motor co-morbidities including sleep and psychiatric disturbances, which are thought to depend on concomitant alterations of serotonergic and noradrenergic transmission. A primary locus of serotonergic neurons is the dorsal raphe nucleus (DRN), providing brain-wide serotonergic input. Here, we identified electrophysiological and morphological parameters to classify serotonergic and dopaminergic neurons in the murine DRN under control conditions and in a PD model, following striatal injection of the catecholamine toxin, 6-hydroxydopamine (6-OHDA). Electrical and morphological properties of both neuronal populations were altered by 6-OHDA. In serotonergic neurons, most changes were reversed when 6-OHDA was injected in combination with desipramine, a noradrenaline reuptake inhibitor, protecting the noradrenergic terminals. Our results show that the depletion of both noradrenaline and dopamine in the 6-OHDA mouse model causes changes in the DRN neural circuitry.

## Introduction

Parkinson’s Disease (PD) is a frequent neurodegenerative disorder characterized by the progressive loss of dopaminergic (DA) neurons in the nigrostriatal pathway, leading to bradykinesia, tremor, rigidity, and postural instability [1, 2]. These cardinal motor symptoms are typically addressed by administration of dopaminergic drugs or by deep brain stimulation. PD patients also experience non-motor symptoms including sleep, affective, and cognitive dysfunctions often preceding the motor disabilities [3, 4]. These comorbidities are in large part refractory to current PD treatments and are thought to be caused by neurodegenerative processes occurring in concomitance to the loss of midbrain dopaminergic neurons. However, the pathology underlying non-motor symptoms remains poorly understood.

Post-mortem studies in PD patients provided first insights into the brain areas which might be involved in the etiology of non-motor dysfunctions in PD. Besides the profound degeneration of the substantia nigra pars compacta (SNc), these studies found cell loss and reduced neurotransmitter release in other monoaminergic brain regions, including the dorsal and median raphe nuclei (DRN and MRN, respectively), and the locus coeruleus (LC) [1, 5-8]. The DRN constitutes the main source of serotonin in the brain with serotonergic cells (DRN^5-HT^) accounting for 30-50% of its neurons [9]. DRN^5-HT^ neurons have been implicated in numerous neuropsychiatric diseases, rendering them a potential neural substrate for non-motor symptoms in PD. DRN^5-HT^ neurons are also of central interest in PD research because of their bidirectional, monosynaptic connection with the striatum [10, 11]. In fact, several studies have shown that serotonergic markers and transmitter levels are altered in Parkinson patients as well as in non-human primate and rodent models of PD [12-19]. Notably, alterations in the serotonergic system have also been related to non-motor comorbidities in PD [20, 21]. Yet, functional investigations of DRN^5-HT^ in rodent models of PD have led to conflicting results showing both increased and decreased activity in DRN^5-HT^ neurons themselves as well as in their downstream targets [22-24]. Besides the serotonergic neurons, the DRN comprises other neuronal populations, including a small group (∼1000 neurons in rats) of DA neurons (DRN^DA^)[25]. DRN^DA^ neurons have been linked to the regulation of pain, motivational processes, incentive memory, wakefulness and sleep-wake transitions [26-30], but their ultimate behavioral significance is yet to be elucidated [31-35]. DRN^DA^ neurons are directly innervated by dopaminergic neurons in the midbrain and have been found to show Lewy bodies in PD patients [6, 30, 36]. Yet, the physiology and pathophysiology of DRN^DA^ neurons in PD remains elusive. The sparsity of research on DRN^DA^ neurons is likely due to the technical challenges associated with targeting this population among the diverse cell types in the DRN and adjacent structures (e.g., retrorubral field, periaqueductal grey and LC), which often co-express signature genes, hampering their molecular identification and region-specific manipulations with cre driver lines [9, 36-39].

Recently, this issue has been addressed by Pinto and colleagues who showed that DRN^DA^ neurons are most faithfully labelled in transgenic mice in which the expression of cre is linked to the DA transporter (DAT-cre) [36]. Previously, the membrane properties of DRN^DA^ neurons have only been addressed in mice in which DRN^DA^ neurons were identified based on the expression of the transcription factor Pitx3 or the enzyme tyrosine hydroxylase (TH) [37]. In Pitx3-GFP mice, about 70% of GFP-positive neurons are TH-positive (TH+) as shown by immunohistochemistry. Moreover, 40% of TH+ neurons in the DRN are not labelled in these mice, suggesting that this line targets a subpopulation of DRN^DA^ neurons [37]. The widely used TH-cre reporter line has been found to show ectopic expression of cre in non-dopaminergic neurons, probably caused by a transient developmental expression of TH [40, 41]. In addition, the TH-cre line also labels noradrenergic neurons in the neighboring LC [40], which produces most of the noradrenaline (NA) in the brain and is involved in mood control, cognition, and sleep regulation [42]. The large overlap of functions ascribed to the LC and DRN is thought to result from the complex reciprocal synaptic connections between these two brain areas: notably, the LC provides noradrenergic input to the DRN [43, 44] while receiving input from DRN^5-HT^ neurons [45-47].

Here, we used ex vivo whole-cell patch clamp recordings and morphological reconstructions to characterize the electrophysiological and morphological properties of DRN^DA^ and DRN^5-HT^ neurons in wild type and DAT-tdTomato mice. Moreover, we studied the impact of catecholamine depletion on DRN^DA^ and DRN^5-HT^ populations in the 6-OHDA toxin model of PD.

## Results

### DRN^DA^ and DRN^5-HT^ neurons are electrophysiologically distinct cell-types

To investigate the electrophysiological and morphological profile of DRN^DA^ neurons and to compare it to DRN^5-HT^ neurons, we performed whole-cell patch clamp recordings in coronal slices of adult wild type and DAT-cre mice crossed with tdTomato reporter mice (Figure 1A). All neurons were filled with neurobiotin and Alexa488 while recording. Alexa488 allowed us to take snapshots of recorded neurons at different time points, thus facilitating the topographical registration of recorded neurons to the post-hoc stained slices (Suppl. Figure 1). Using this approach, we obtained complete sets of electrophysiological and morphological data from 75 neurons in the DRN. Cells were identified as DRN^5-HT^ or DRN^DA^ neurons based on tryptophan hydroxylase (TPH) or TH immunoreactivity, respectively (Figure 1A-D). In line with Fu et al. (2010)[38], none of the recorded neurons was positive for both TPH and TH (n = 0/412). During the recordings, we used a series of depolarizing and hyperpolarizing current steps and ramps that allowed us to characterize active and passive membrane properties in detail (Figure 1E – G). Based on the electrophysiological data, we first tested possible differences between TH+ neurons recorded in wild-type mice and tdTomato-positive (tdTomato+) neurons recorded in DAT-tdTomato mice. We found no differences between these two groups (n = 13 TH+ vs n = 30 tdTomato+ neurons, Suppl. Figure 2A) and neither within the subset of tdTomato+ neurons when comparing TH+ to TH-negative (TH-) neurons (n = 23 TH+ vs n = 6 TH-neurons, Suppl. Figure 2A–E). Since this small number of TH-neurons were positive for DAT and their electrophysiology indistinguishable from TH+ DRN^DA^ neurons, the data was pooled. Please note that staining of recorded neurons, i.e. immunohistochemistry on slices strained by hour-long patch-clamp recordings, is more challenging as neurons can be lost after patching (no staining data) or the staining might be ambiguous. Out of 114 tdTomato+ neurons only one cell displayed a different electrophysiological profile than all other DRN^DA^ neurons, suggesting a false-positive rate of 0.8%. That neuron was TH-,displayed profoundly distinct intrinsic properties, and was therefore excluded (Suppl. Figure 2F, G). Taken together, the electrophysiological results support the use of the DAT-tdTomato mouse line when studying DRN^DA^ neurons and data from both mouse lines were pooled. Recordings of DRN^DA^ neurons revealed distinctive electrophysiological properties such as a slowly ramping membrane potential during constant current injections giving rise to delayed spiking and postinhibitory hypoexcitability (Figure 1E). Moreover, most DRN^DA^ neurons displayed rebound oscillations and sag currents (Figure 1E, F).

**Figure 1.**
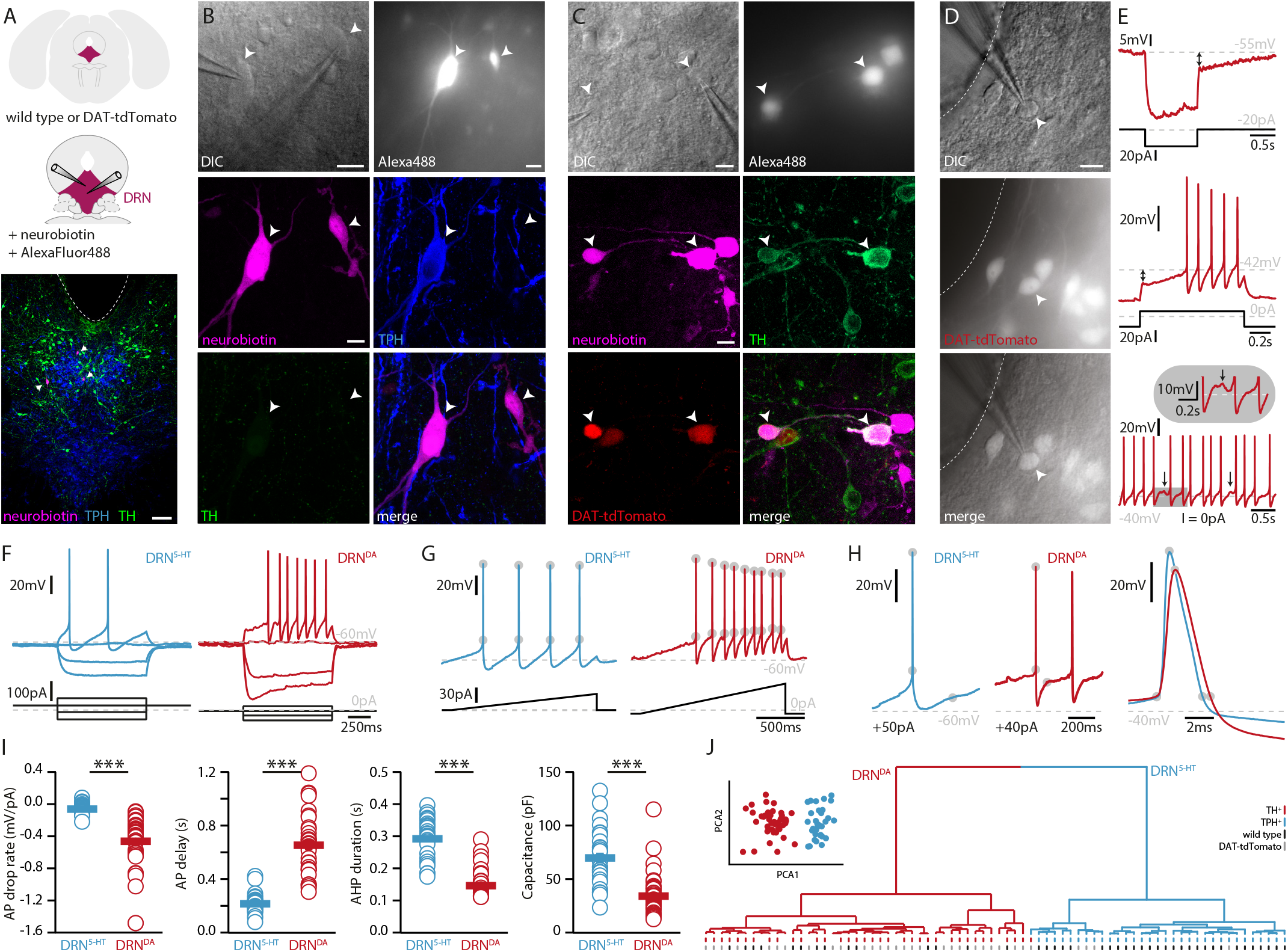
DRN^DA^ and DRN^5-HT^ are electrophysiologically distinct cell-types. **(A)** Scheme of the location of the DRN (pink) in a coronal section (top) and at higher magnification together with two patch pipettes (center). Bottom: a representative slice stained post recording for TPH, TH and neurobiotin revealing serotonergic neurons (arrows). The ventricle is indicated with a dashed line. **(B)** Top: differential interference contrast (DIC) microscopy image (left) of neurons that were filled with Alexa488 (right) and neurobiotin. Center, bottom: staining of the same neurons revealing a TPH+ (DRN^5-HT^) neuron and a TPH- and TH-cell. **(C)** Top: DIC image of recorded neurons that were filled with Alexa488 and neurobiotin. Center, bottom: staining of the same neurons revealing tdTomato- and TH+ (DRN^DA^) neurons. **(D)** Representative fluorescent (top), DIC (center) image and overlay (bottom) of a tdTomato+ neuron in a DAT-tdTomato mouse. **(E)** Representative recordings depicting postinhibitory hypoexcitability, slowly ramping currents and rebound oscillations in DRN^DA^ neurons. **(F)** Representative voltage responses to current injections in a TPH+ (DRN^5-HT^) and TH+ (DRN^DA^) neuron. **(G)** Ramping current injections reveal action potential (AP) amplitude accommodation. Gray circles indicate the onset and peak of APs. **(H)** Amplitude and duration of the AP and AP afterhyperpolarization (AHP) in a DRN^5-HT^ and DRN^DA^ neurons. Gray circles indicate onset, peak, and end of the AP and AHP. **(I)** Quantification of electrophysiological properties distinguishing DRN^5-HT^ from DRN^DA^ neurons (AP drop rate: n = 32 DRN^5-HT^, n = 43 DRN^DA^, Capacitance: n = 32 DRN^5-HT^, n = 43 DRN^DA^, AP delay: n = 30 DRN^5-HT^, n = 43 DRN^DA^, AHP duration: n = 28 DRN^5-HT^, n = 32 DRN^DA^, N = 9; Wilcoxon Rank Sum Test). **(J)** Principal component analysis (PCA) of five electrophysiological parameters (insert) and hierarchical cluster analysis based on PCA1 and PCA2 (Ward’s method, Euclidean distance). Intrinsic properties were sufficient to separate TPH+ cells (blue dash) from TH+ (red dash) cells. Bottom dashes indicate wild type (black) and DAT-tdTomato (gray) mice. Data are shown as mean ± SEM, ***p < 0.001. Scale bars: A, 100 μm; B-D, 10 μm.

**Figure 2.**
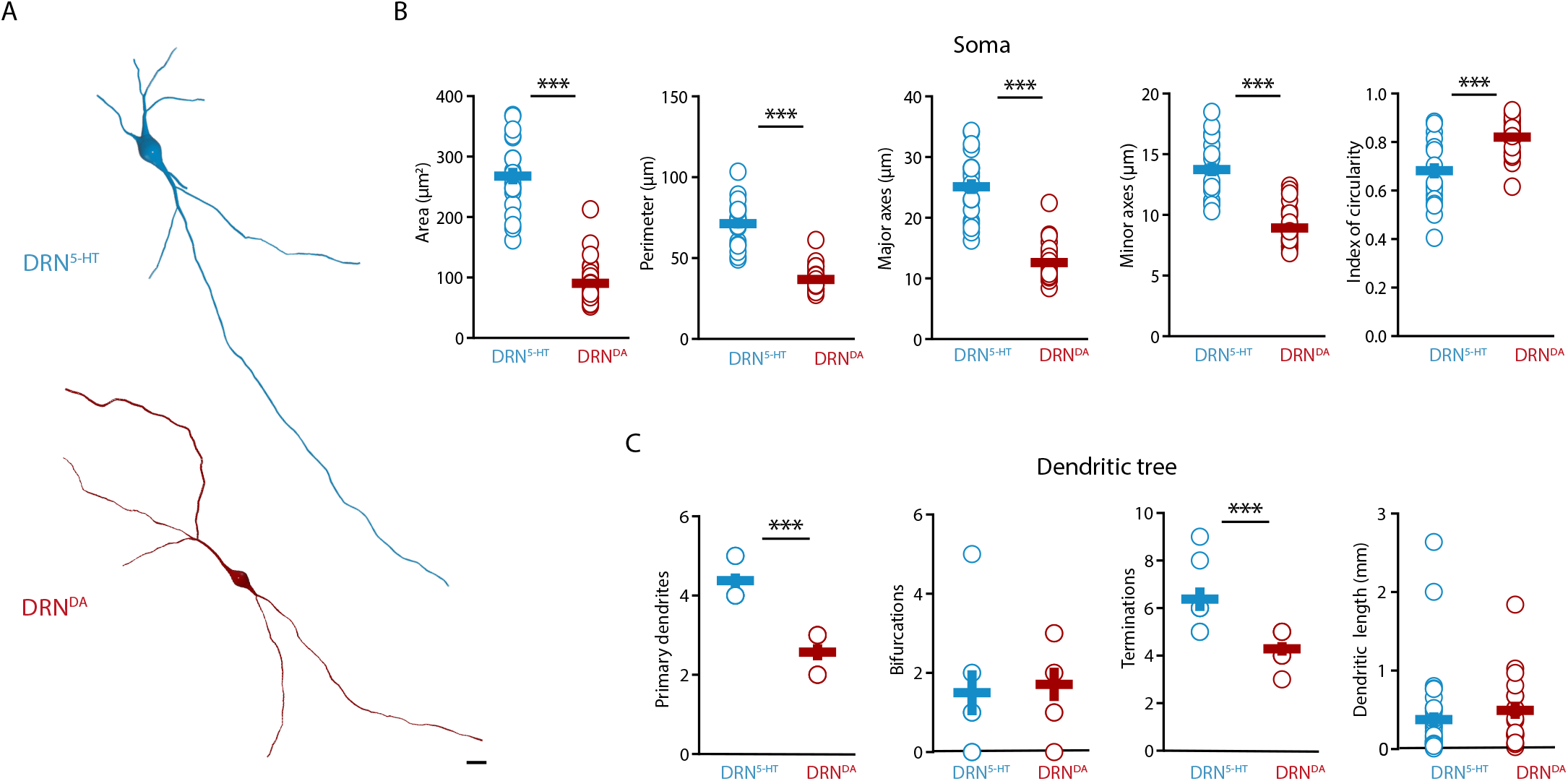
DRN^DA^ and DRN^5-HT^ have distinct morphological profiles. **(A)** Top: representative digital reconstruction of a DRN^5-HT^. Bottom: representative digital reconstruction of a DRN^DA^. **(B-F)** Morphological parameters describing the soma size and shape of DRN^5-HT^ and DRN^DA^ neurons (DRN^5-HT^: n = 20, N = 3; DRN^DA^: n = 27, N = 3; unpaired t-test or Mann-Whitney U test). **(G-J)** Morphological parameters describing the dendritic tree of DRN^5-HT^ and DRN^DA^ neurons (DRN^5-HT^: n = 8, N = 3; DRN^DA^: n = 7, N = 3; Mann-Whitney U test). Data are shown as mean ± SEM, ***p < 0.001. Scale bar: 10 μm.

When comparing the electrophysiological properties of DRN^DA^ to DRN^5-HT^ neurons, we observed numerous differences between these two cell types, but here we focus on the five most significant ones. While DRN^5-HT^ neurons spike with short delays in response to current steps and maintain a relatively constant action potential (AP) amplitude, DRN^DA^ neurons display a longer delay to the first spike and the amplitude of subsequent APs drops (Figure 1F-I). Additionally, the APs of DRN^5-HT^ neurons rise faster, while their afterhyperpolarization (AHP) is longer compared to DRN^DA^ neurons (Figure 1H, I). Lastly, the capacitance of DRN^5-HT^ neurons is significantly larger than in DRN^DA^ neurons (Figure 1I).

Next, we tested if DRN^DA^ neurons can be distinguished from DRN^5-HT^ neurons based on these five electrophysiological parameters. To this end, we standardized the data and ran a principal component analysis (PCA) including all DRN^5-HT^ neurons (i. e. all TPH-positive, TPH+), all TH+ neurons recorded in wild-type mice and all tdTomato+ cells recorded in DAT-tdTomato mice (except for one outlier shown in Suppl. Figure 2F, G). Plotting the first two principal components (PC) showed two separate clusters (Figure 1J, insert). Unsupervised hierarchical cluster analysis based on PC1 and PC2 revealed the same two major clusters and potential subclusters (Figure 1J). Mapping the molecular identity of the cells onto the dendrogram revealed the separation of DRN^5-HT^ and DRN^DA^ neurons, while there was no branching according to mouse line (wild type vs. DAT-tdTomato), further corroborating the validity of DAT-tdTomato mice as a marker for DRN^DA^ neurons. Overall, these data suggest that electrophysiological parameters themselves are sufficient to distinguish between DRN^5-HT^ and DRN^DA^ neurons.

In addition to DRN^DA^ and DRN^5-HT^ the DRN contains an unknown number of cell types and 47 out of 120 recorded neurons were neither TH+, nor TPH+ and did not express tdTomato. To test whether DRN^DA^ can also be distinguished from those populations based on their electrophysiological profile, we ran a PCA on 20 standardized parameters and used the first three PCAs for unsupervised hierarchical clustering (Suppl. Figure 3). Our analysis suggests that there might be four major electrophysiological cell types in the DRN. In contrast to DRN^DA^ and DRN^5-HT^ neurons, a large proportion of the remaining cells showed rebound spiking and biphasic AHPs, resembling the profiles of local interneurons in other brain areas (Suppl. Figure 3D, E). Interestingly, the clustering also indicated that three TH- and tdTomato-negative (tdTomato-) neurons belonged to the DRN^DA^ neurons and further analysis showed that they were indistinguishable from molecularly identified DRN^DA^ neurons (Suppl. Figure 3C, F). These findings indicate that clustering can be used to identify neurons that otherwise would have been excluded due to a lack of post-hoc staining data or genetic driver lines.

**Figure 3.**
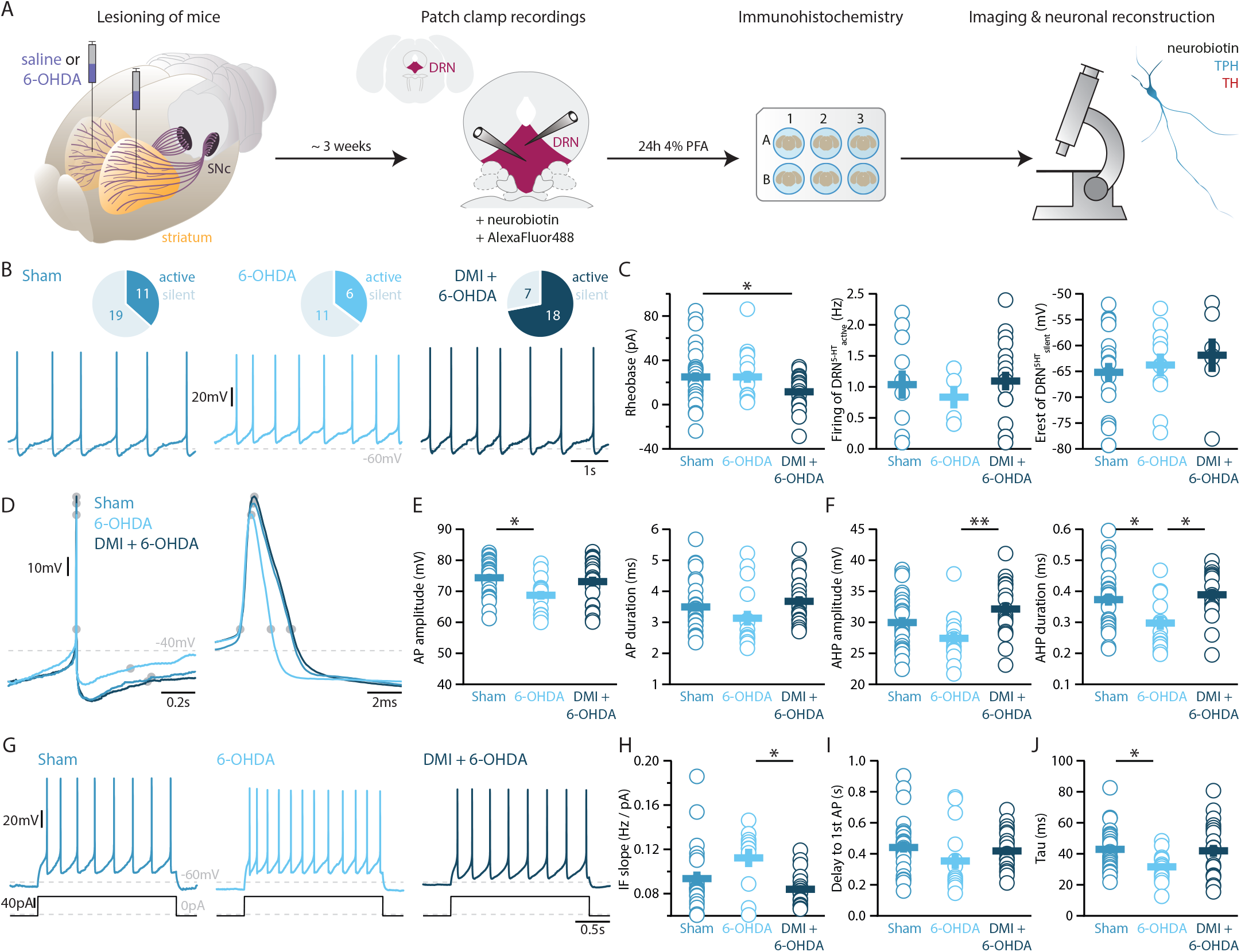
Lesions targeting primarily nigrostriatal dopamine increase the excitability of DRN^5-HT^ neurons whereas loss of NA affects their action potentials. **(A)** Overview of workflow for analyzing the electrophysiological and morphological properties of DRN neurons in Sham and 6-OHDA lesioned mice. **(B)** Top: pie charts showing the number of spontaneously active (dark) and silent (pale) DRN^5-HT^ neurons in three conditions: Sham (left), 6-OHDA-injected mice (center) and 6-OHDA-injected mice pre-treated with desipramine (DMI + 6-OHDA, right). Bottom: representative recordings of spontaneously active DRN^5-HT^ neurons (I = 0 pA). (C) Quantification of the rheobase (left, Sham: n = 30, 6-OHDA: n = 17, DMI + 6-OHDA: n = 25), the firing frequency of spontaneously active cells (center, Sham: n = 11, 6-OHDA: n = 6, DMI + 6-OHDA: n = 18), and the resting membrane potential of silent DRN^5-HT^ neurons (right, Sham: n = 19, 6-OHDA: n = 11, DMI + 6-OHDA: n = 7). **(D)** Representative action potentials of DRN^5-HT^ at low (left) and high (right) temporal resolution. Gray circles indicate onset, offset, and peak of the APs as well as the end of the afterhyperpolarization (AHP). **(E)** Quantification of the amplitude (left) and duration (right) of the APs of DRN^5-HT^ neurons (Sham: n = 29, 6-OHDA: n = 16, DMI + 6-OHDA: n = 21). **(F)** Same as in **(D)** for the AHP. (G) Representative responses of DRN^5-HT^ neurons to current steps (I = +75 pA). **(H)** Quantification of firing frequency / injected current. **(I)** Quantification of the delay to the first AP when injected with current eliciting 1Hz firing (Sham: n = 29, 6-OHDA: n = 16, DMI + 6-OHDA: n = 21). **(J)** Quantification of the membrane time constant (tau) of DRN^5-HT^ neurons (Sham: n = 32, 6-OHDA: n = 16, DMI + 6-OHDA: n = 21). Sham: N = 6 - 7; 6-OHDA: N = 7; DMI + 6-OHDA: N = 4; unpaired t-test or Mann-Whitney U test. Data are shown as mean ± SEM, * p < 0.05, ** p < 0.01.

Overall, our data show that DRN^DA^ neurons constitute an electrophysiologically distinct class of neurons in the DRN expressing several hallmark properties, which are sufficient to identify them within the local DRN circuitry.

### DRN^DA^ and DRN^5-HT^ neurons have different morphological properties

Next, we characterized the morphological profile of DRN^5-HT^ and DRN^DA^ neurons. We focused on the analysis of somatic and dendritic properties since a complete reconstruction of the axonal arborization could not be retrieved from the slices. The analysis of the somatic properties showed that DRN^5-HT^ neurons had larger cell bodies than DRN^DA^ neurons (Figure 2A, B), as measured in their area, perimeter, length, and width (Figure 2B). Cell bodies also differed in shape, with DRN^DA^ neurons having more circular somata than DRN^5-HT^ neurons, as indicated by the circularity index (Figure 2B). Analyzing the dendritic properties, we found that DRN^5-HT^ neurons had four to five primary dendrites, compared to only two to three in DRN^DA^ neurons (Figure 2A, C). Moreover, dendrites of DRN^DA^ neurons were frequently bipolar with the main primary dendrites starting from opposite extremes of the soma. Both populations had relatively few bifurcations (Figure 2C), but the DRN^5-HT^ neurons had significantly more terminations (Figure 2C). The overall dendritic length did not differ between the DRN^5-HT^ and DRN^DA^ neurons: both populations had a mix of short and long dendrites (Figure 2C). These data suggest that DRN^5-HT^ neurons have denser dendritic arborization than DRN^DA^ neurons, mostly due to larger numbers of primary dendrites.

Altogether, our results show that DRN^5-HT^ and DRN^DA^ neurons have distinct morphological properties. DRN^5-HT^ neurons are mostly multipolar neurons, with a big and complex soma and multiple primary dendrites, while DRN^DA^ neurons have smaller and more circular cell bodies with bipolar dendrites.

### DA and NA depletion distinctly affect the membrane properties of DRN^5-HT^ neurons

To elucidate how DRN^5-HT^ and DRN^DA^ neurons might be affected in PD, we characterized these populations in a mouse model of PD based on bilateral injection of the neurotoxin 6-OHDA in the dorsal striatum. This approach leads to a partial lesion of catecholamine neurons, reproducing an early stage of parkinsonism in which particularly non-motor symptoms such as depression- and anxiety-like behavior are manifested [48, 49]. In line with previous studies, we observed a 60-70% reduction of TH levels in the striatum (Suppl. Figure 4, [50]). Only mice meeting this criterion were included in the study. Measurement performed by ELISA showed that the 6-OHDA injection did not alter the levels of 5-HT in the striatum (Suppl. Figure 4D), and immunostaining showed that the striatal 6-OHDA injection did not cause degeneration of DRN^5-HT^ or DRN^DA^ neurons (Suppl. Figure 5).

**Figure 4.**
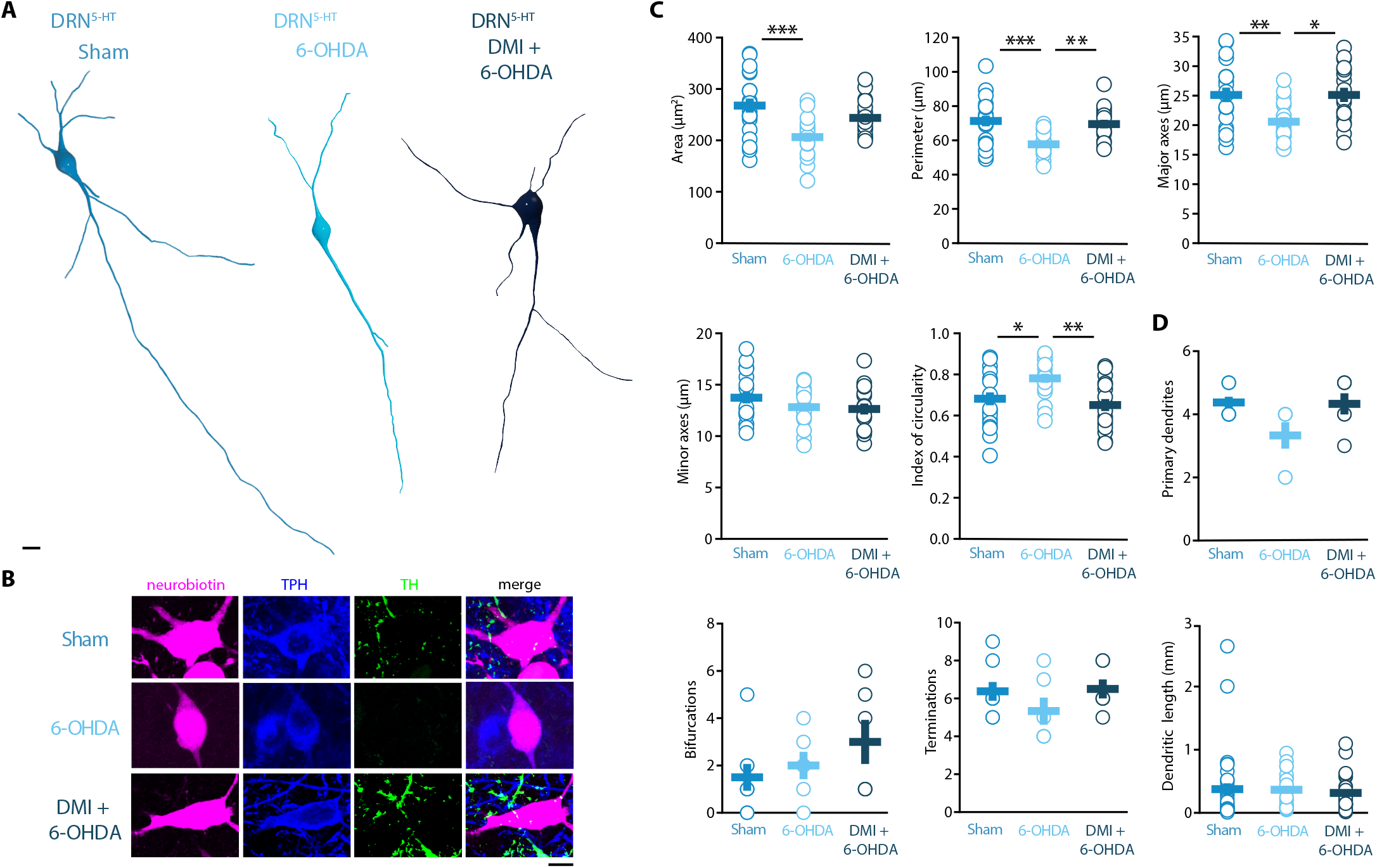
Striatal injection of 6-OHDA induced a hypotrophic phenotype in the DRN^5-HT^, which is prevented by pre-treatment with DMI. **(A)** Representative digital reconstructions of a DRN^5-HT^ neuron in three different conditions: Sham (left), 6-OHDA-injected mice (center) and 6-OHDA-injected mice pre-treated with desipramine (right). **(B)** Representative confocal pictures of soma from DRN^5-HT^ neurons in Sham (top), 6-OHDA-injected mice (center), and 6-OHDA-injected mice pre-treated with desipramine (bottom). **(C-G)** Morphological descriptors of the soma size and shape in DRN^5-HT^ neurons (Sham: n = 20, N = 4; 6-OHDA: n = 19, N = 4; DMI + 6-OHDA: n = 17, N = 3; one-way ANOVA). **(H-K)** Morphological descriptors of the dendritic tree in DRN^5-HT^ neurons (Sham: n = 8, N = 3, 6-OHDA: n = 6, N = 3: DMI + 6-OHDA: n = 6, N = 2). Data are shown as mean ± SEM, ***p<0.001, **p<0.01, *p<0.05. Scale bar: 10 μm.

**Figure 5.**
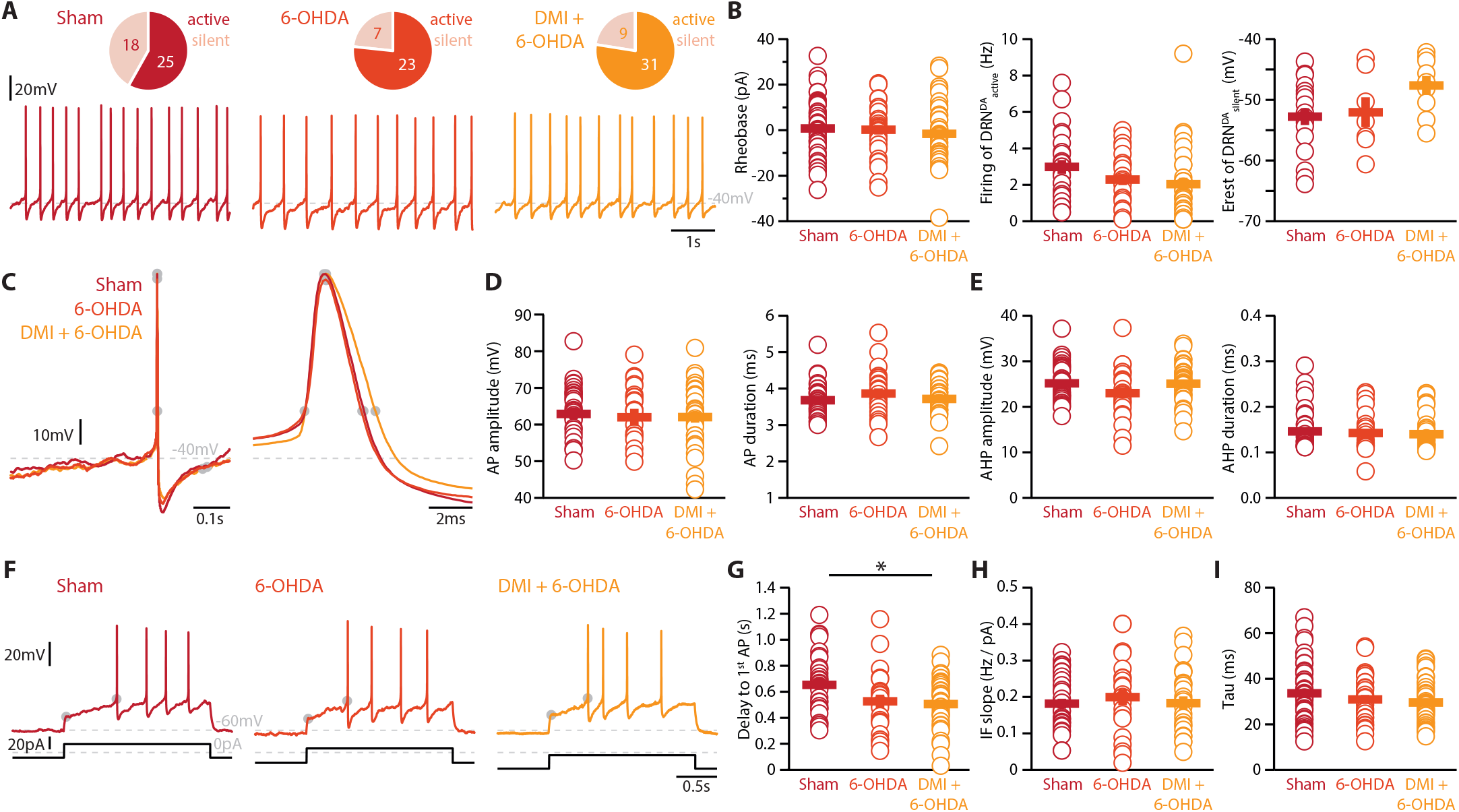
Lesions targeting primarily SN dopamine depolarize DRN^DA^ neurons whereas concomitant loss of NA does not affect their action potentials. **(A)** Top: Pie charts showing the proportion of spontaneously active (dark) and silent (pale) DRN^DA^ neurons in three conditions: Sham (left), 6OHDA-injected mice (center) and 6-OHDA-injected mice pre-treated desipramine (DMI, right). Bottom: Representative recordings of spontaneously active DRN^DA^ (I = 0pA). **(B)** Quantification of the rheobase (left, Sham: n = 43, 6-OHDA: n = 31, DMI + 6-OHDA: n = 40), the firing frequency of spontaneously active (center, Sham: n = 25, 6-OHDA: n = 23, DMI + 6-OHDA: n = 31), and the resting membrane potential of silent DRN^DA^ neurons (right, Sham: n = 18, 6-OHDA: n = 7, DMI + 6-OHDA: n = 9). **(C)** Representative action potentials of DRN^DA^ at low (left) and high (right) temporal resolution. Gray circles indicate onset, offset, and peak of APs and the end of the afterhyperpolarization (AHP). **(D)** Quantification of the amplitude (left) and duration (right) of the APs of DRN^DA^ neurons (Sham: n = 34, 6-OHDA: n = 23, DMI + 6-OHDA: n = 35). **(E)** Same as in **(D)** for the AHP. **(F)** Representative responses of DRN^DA^ neurons to current steps (I = 75pA). Gray circles indicate the delay to the first AP. **(G-I)** Quantification of firing frequency / injected current (G, Sham: n = 31, 6-OHDA: n = 23, DMI + 6-OHDA: n = 27), the delay to the first AP when injected with current eliciting 2 Hz firing (H, Sham: n = 34, 6-OHDA: n = 23, DMI + 6-OHDA: n = 35), and the membrane time constant (I, Sham: n = 43, 6-OHDA: n = 29, DMI + 6-OHDA: n = 40) of DRN^DA^ neurons recorded (Sham: N = 8; 6-OHDA: N = 6; DMI + 6-OHDA: N = 6; unpaired t-test or Mann-Whitney U test). Data are shown as mean ± SEM, * p < 0.05.

Striatal injection of 6-OHDA has also been found to produce a partial loss of NA neurons in the LC [50] and ELISA analysis showed that this approach induces approximately 60% loss of NA in the striatum (Suppl. Figure 4E). In the present study, we determined the specific impact of NA dysfunction on the physiology of DRN^5-HT^ and DRN^DA^ neurons by pre-treating a group of mice with desipramine (DMI), a selective inhibitor of NA reuptake, before injecting 6-OHDA (DMI+6-OHDA mice), which partially prevents striatal NA loss (Suppl. Figure 4E). We then assessed the intrinsic properties of DRN^5-HT^ and DRN^DA^ neurons in Sham-lesion (Sham), 6-OHDA- and DMI+6-OHDA-treated mice (Figure 3A). Whole-cell recordings obtained from DRN^5-HT^ neurons in control mice revealed that 37% of DRN^5-HT^ neurons were spontaneously active in slices and the proportion of intrinsically active neurons was similar in mice injected with 6-OHDA (Sham: n = 11/30 DRN^5-HT^ neurons, 6-OHDA: n = 6/17 DRN^5-HT^ neurons, Figure 3B). However, DRN^5-HT^ neurons recorded in DMI+6-OHDA mice showed an increased excitability: in this condition, 72% of DRN^5-HT^ neurons were spontaneously active and DRN^5-HT^ neurons displayed lower rheobase currents than control mice (Figure 3B, C). Because of the protective effect exerted in these mice by DMI, these findings suggest that the p noradrenergic system contributes to the increased firing of DRN^5-HT^ neurons.

While the rheobase of DRN^5-HT^ neurons was not affected in 6-OHDA mice, we observed that their firing properties were profoundly altered: DRN^5-HT^ neurons recorded in 6-OHDA mice displayed smaller APs than Sham mice and shorter AHPs than both Sham and 6-OHDA injected mice pre-treated with DMI (Figure 3D-F). In contrast, the APs and AHPs of Sham and 6-OHDA injected mice pretreated with DMI did not differ. Moreover, DRN^5-HT^ neurons of 6-OHDA injected mice fired at higher frequencies than 6-OHDA injected mice pretreated with DMI (Figure 3G-I). Finally, the membrane time constant of DRN^5-HT^ neurons was shorter in 6-OHDA injected mice than in Sham mice (Figure 3J). Interestingly, we found no differences in the firing properties of DRN^5-HT^ neurons recorded in Sham and in 6-OHDA injected mice pre-treated with DMI, suggesting that the noradrenergic lesion critically contributes to the changes in 6-OHDA mice. Taken together, these results indicate that DRN^5-HT^ neurons are affected in the 6-OHDA mouse model of PD. Specifically, lesions of the dopaminergic system increase the excitability of DRN^5-HT^ neurons whereas the combined lesion of the noradrenergic and dopaminergic systems changes the firing properties of DRN^5-HT^ neurons.

### Striatal DA depletion induces hypotrophy of DRN^5-HT^ neurons

Morphological analysis revealed a reduced soma size of the DRN^5-HT^ neurons in 6-OHDA mice, which was manifested as decreased area, perimeter, and major axes in comparison to control mice (Figure 4A-C). Moreover, the increase in the circularity of the 6-OHDA group indicated that the shape of the soma of DRN^5-HT^ neurons was also altered by the lesion (Figure 4C). These modifications were not observed in DMI + 6-OHDA mice, suggesting that preserving the NA system protected the DRN^5-HT^ neurons (Figure4A-C). Finally, the injection of 6-OHDA without DMI pre-treatment also resulted in a trend toward reduced number of primary dendrites and terminations of DRN^5-HT^ neurons (Figure 4D). The number of bifurcations and the dendritic length were not affected by the lesion (Figure 4D). Globally, these results suggest that the lesion produced by 6-OHDA induces a hypotrophic phenotype in DRN^5-HT^ neurons characterized by a shrinkage of the soma and that this alteration is NA-dependent.

### Striatal DA depletion affects the firing of DRN^DA^ neurons independent of NA loss

Finally, we assessed whether the striatal 6-OHDA lesion affects the physiology of DRN^DA^ neurons. Whole-cell patch-clamp recordings revealed that 58% of DRN^DA^ neurons are spontaneously active in slices of Sham-lesion mice (Figure 5A). In contrast, the proportion of intrinsically active neurons increased to 77% and 78% of DRN^DA^ neurons in 6-OHDA injected mice with and without pre-treatment with DMI, respectively.

In stark contrast to DRN^5-HT^ neurons, the rheobase, the APs and their AHPs, the IF slope and the time constant of DRN^DA^ neurons were not affected in any 6-OHDA mice (Figure 5B-I). In fact, we did not observe any change in the firing properties of DRN^DA^ neurons that was dependent on the protection of the NA system with DMI (Figure 5). DRN^DA^ neurons recorded in 6-OHDA injected mice pre-treated with DMI did however display a reduction in spike latency compared to Sham-lesioned mice (Figure 5G). Together, these results suggest that the electrophysiological properties of DRN^DA^ neurons are affected in the 6-OHDA mouse model of PD and that these changes are primarily due to the lesion of the nigrostriatal dopaminergic pathway.

### Striatal DA depletion does not change significantly the morphology of DRN^DA^ neurons

The morphological analysis of DRN^DA^ neurons revealed that the striatal 6-OHDA injection did not significantly affect somatic and dendritic morphology.

### Unilateral lesion of LC NA cells induces minor changes in DRN subpopulations

Our results so far suggest that concomitant lesioning of the DA and NA system (6-OHDA model) has a severe impact on DRN^5-HT^ neurons which cannot be evoked when the NA system is partially protected (DMI + 6-OHDA model). Therefore, we next assessed if selective lesioning of the NA system itself is sufficient to evoke changes in electrical and morphological properties observed in DRN^5-HT^ neurons recorded in 6-OHDA. To that end we performed unilateral injections of 6-OHDA/saline in the LC (Suppl. Figure 6A), which lead to approximately 50% loss of TH+ neurons in the LC (Suppl. Figure 6B, C). We chose to restrict the injection of 6-OHDA to one hemisphere because little is known about this type of lesion while the fundamental role of the NA system in various neural processes is well established [51, 52]. We found that selective lesioning of the LC (‘6-OHDA-LC’) did not alter the baseline activity levels, firing frequencies and resting membrane potentials of DRN^5-HT^ and DRN^DA^ neurons (Suppl. Fig. 6D, E, K, L). However, DRN^5-HT^ neurons recorded in 6-OHDA-LC had a lower input resistance at hyperpolarized membrane potentials, a shorter afterhyperpolarization and a larger capacitance than DRN^5-HT^ neurons recorded in control mice (Sham-LC, Suppl. Fig. 6E-G). Moreover, DRN^DA^ neurons recorded in 6-OHDA-LC showed a reduction in their sag amplitudes (Suppl. Fig 6M, N). The other electrophysiological parameters were not significantly affected. Morphological analysis revealed that the selective lesion of the noradrenergic system did not alter the size and shape of cell bodies in either DRN subpopulation (Suppl. Figure 6H,I, O,P), however, the dendritic branching of both subpopulations was altered, as shown by the increased length of primary dendrites in DRN^5-HT^neurons (Suppl. Figure 6J) and the increased number of primary dendrites in DRN^DA^ neurons (Suppl. Figure 6Q).

**Figure 6.**
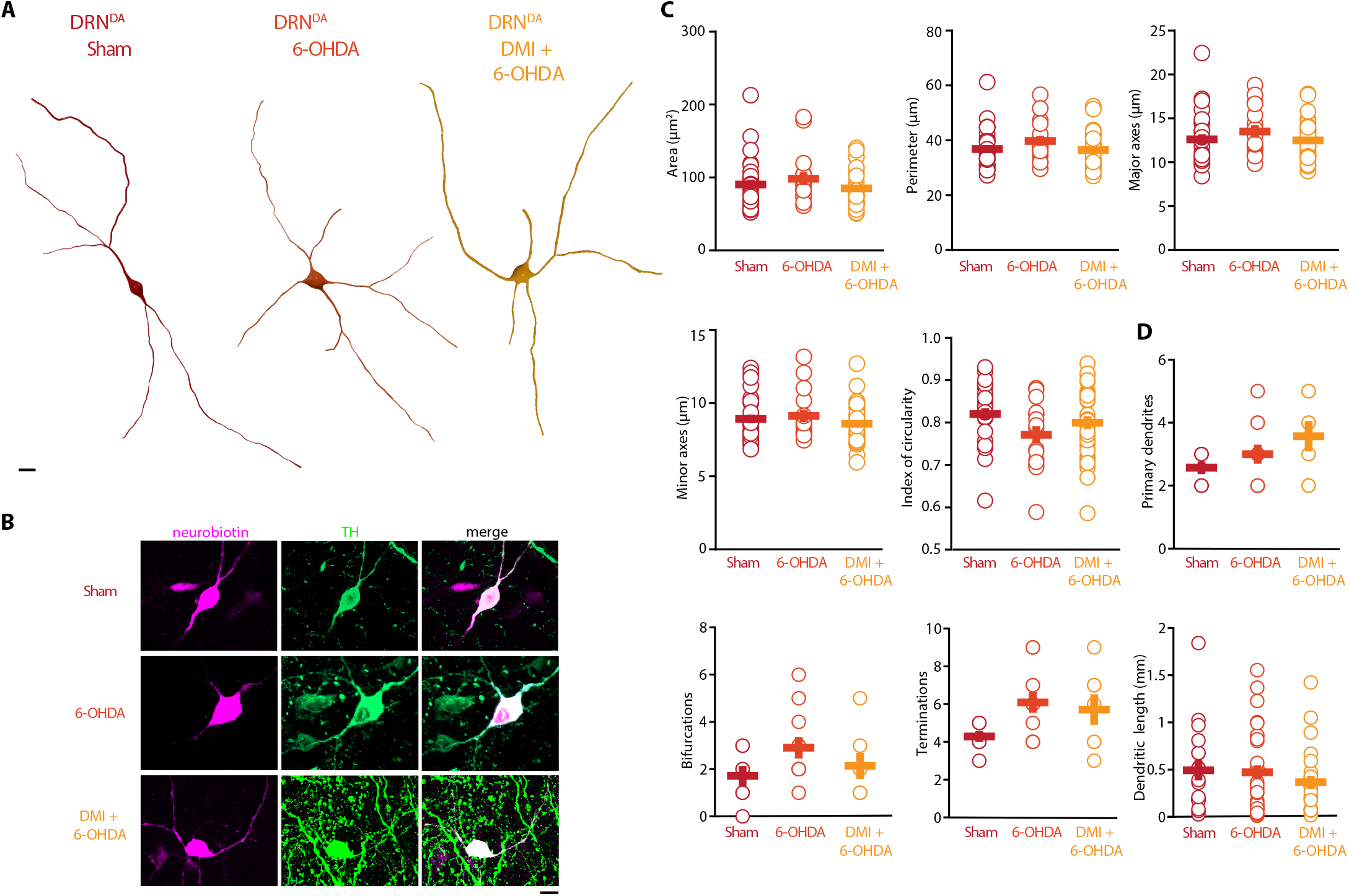
Striatal injection of 6-OHDA did not alter morphology of DRN^DA^. **(A)** Representative digital reconstructions of a DRN^DA^ neuron in three different conditions: Sham (left), 6-OHDA-injected mice (center) and 6-OHDA-injected mice pre-treated with desipramine (right). **(B)** Representative confocal pictures of soma from DRN^DA^ neurons in Sham (top), 6-OHDA-injected mice (center), and 6-OHDA-injected mice pre-treated with DMI (bottom). **(C-G)** Morphological descriptors of the soma size and shape in DRN^DA^ neurons (Sham: n = 27, N = 7, 6-OHDA: n = 16, N = 4; DMI + 6-OHDA: n = 31, N = 5). **(H-K)** Morphological descriptors of the dendritic tree in DRN^DA^ neurons. (Sham: n = 7, N = 3; 6-OHDA: n = 11, N = 4; DMI + 6-OHDA: n = 7, N = 3). Data are shown as mean ± SEM. Scale bar: 10 μm.

## Discussion

In the present study we combine *ex vivo* whole-cell patch clamp recordings with morphological reconstructions and immunohistochemistry, to show that DRN^DA^ neurons have a distinct electrophysiological profile, which is sufficient to distinguish them from DRN^5-HT^ neurons as well as other neuron classes in the DRN. Utilizing this approach, we also reveal that, in a 6-OHDA mouse model of PD, DRN^5-HT^ neurons display distinct pathophysiological changes depending on the loss of DA and NA. Notably, degeneration of noradrenergic neurons affects not only the electrical properties of DRN^5-HT^ neurons but also evokes hypotrophy of their cell bodies. In contrast, the loss of nigrostriatal DA mainly affects the electrophysiological properties of DRN^DA^ neurons while concomitant loss of NA alters their morphology.

We used an extensive electrophysiological characterization protocol to quantify the differences between the DRN^DA^ and DRN^5-HT^ populations. The electrophysiological properties agree with previous studies, such as the spontaneous firing pattern seen in DRN^DA^ neurons [37], and the slow AHP of DRN^5-HT^ neurons [23]. Standard electrophysiological parameters were used to create a classification tool, which efficiently identifies DRN^5-HT^ and DRN^DA^ cells, including DA neurons confirmed by TH staining and / or by fluorescent expression in DAT-tomato mice (Figure 1). Importantly, the DRN^DA^ neurons recorded from wild-type and DAT-tdTomato mice did not differ in their electrical properties, indicating that the transgene does not interfere with the membrane properties of this population.

We showed that DRN^DA^ neurons share electrophysiological properties with other dopaminergic populations in the midbrain such as postinhibitory hypoexcitability, rebound oscillations, a slowly ramping membrane potential and sag currents [37, 53, 54], yet their electrophysiological profile is distinct from DRN^5-HT^ neurons as well as other neuronal populations in the DRN. Most of the parameters extracted in our characterization rely on intracellular recordings of the membrane potential. However, some properties such as spontaneous firing and AP kinetics could be useful for *in vivo* characterization, even in extracellular recordings [55-57]. In addition to the DRN^5-HT^ and DRN^DA^ neuronal populations, a large fraction of neurons displayed electrophysiological properties that were distinct from these two groups (Suppl. Figure 3), suggesting that there are other neuronal subtypes in the DRN network, such as previously reported GABAergic, glutamatergic, and peptidergic neurons [9, 10, 58-62].

In line with previous studies, the majority of DRN^5-HT^ neurons were large multipolar or fusiform neurons with four to five primary dendrites, very distinct from the DRN^DA^ neurons [63-65]. Very little is known about the morphology of the DRN^DA^ neurons, but previous studies identified small ovoid cells in the DRN which are likely to correspond to the DRN^DA^ cells [37, 66]. Out of 25 reconstructed DRN^5-HT^ neurons, only one displayed dendritic spines. Previous studies in rats described the presence of dendritic spines in most DRN^5-HT^ neurons [67]. However, the study was performed in thicker slices and the dendritic spines were scarce in the primary and secondary dendrites, while they became dense in the distal dendrites, thus it is possible that in our study those dendrites were not present [67].

In the present study we assessed the impact on DRN cells of a striatal bilateral 6-OHDA lesion performed with or without DMI pre-treatment, which has been shown to protect the NA neurons in the LC from the 6-OHDA-induced degeneration [50, 68, 69]. We found that both DRN^5-HT^ and DRN^DA^ populations were affected in a cell-type specific manner by the combined action of 6-OHDA on DA and NA, with DRN^5-HT^ neurons being particularly sensitive to changes in the noradrenergic system. Loss of SNc dopaminergic neurons alone (6-OHDA + DMI) – which are known to target DRN^5-HT^ and DRN^DA^ neurons directly – increased the excitability and spontaneous activity in DRN^5-HT^ neurons [10]. This is in line with previous *ex vivo* and *in vivo* studies showing that the DRN^5-HT^ neurons display increased firing rates in rodents pre-treated with DMI and injected with 6-OHDA [23, 70] As hypothesized by Prinz et al. [23], the selective loss of midbrain DA may induce a homeostatic increase in the excitability of DRN^5-HT^ neurons. Our data contrast with a previous *in vivo* study showing decreased firing activity in DRN^5-HT^ neurons where injection of 6-OHDA was preceded by treatment with DMI and fluoxetine [24]. This dissimilarity may be related to species-specific (rat vs mouse) and technical (intracerebroventricular vs striatal injections, recordings performed at 10 days vs three weeks after the 6-OHDA injection). Importantly, in the same study identification of DRN^5-HT^ neurons was not molecularly confirmed and the data may include other spontaneously active DRN neurons. In fact, our recordings show that there are non-serotonergic neurons in the DRN, which are spontaneously active and display a regular, slow firing frequency similar to DRN^5-HT^ neurons, highlighting the importance of unequivocal identification of DRN cell types.

The present study shows that combined DA and NA lesioning affects DRN^5-HT^ neurons more profoundly than selective loss of DA (Figure 3, 4). In mice treated with 6-OHDA only, several electrophysiological and morphological properties were altered (Figure 3, 4). The time constant and AHP of DRN^5-HT^ neurons were shorter and the neurons responded with higher firing frequencies to current injections than in Sham. This finding suggests that the pronounced AHP and long tau of these neurons may act as a “brake” limiting their maximum firing frequency in control conditions and that this brake is reduced when the NA system is lesioned. Future studies are needed to assess if DRN^5-HT^ neurons in fact fire at higher rates in vivo in mice treated with 6-OHDA. In contrast, such changes in DRN^5-HT^ neurons were prevented when the NA system was protected by pre-treatment with DMI. These findings indicate an important role for NA as mediator of changes in the activity and properties of DRN^5-HT^ neurons. The changes produced by the 6-OHDA lesion on the DRN^DA^ population were less pronounced than and different from those in DRN^5-HT^ neurons. In terms of electrophysiological properties, the observed changes were primarily in the DA only lesion (6-OHDA + DMI), suggesting that unlike DRN^5-HT^, DRN^DA^ neurons are affected by the loss of midbrain DA rather than the accompanying changes in NA (Figure 5). Interestingly, unilateral lesions in the LC did not result in significant alterations in DRN neurons to the extent of the larger striatal lesions (Suppl. Figure 6). Although the trends of some of the electrophysiological parameters, such as the amplitude and duration of the AP and the AHP observed in DRN^5-HT^ neurons, were similar to those induced by the 6-OHDA only lesion shown in Figure 3, the effects were smaller. This could be due to the more limited extent of the LC injections compared to the striatal ones as well as the unilateral LC vs. bilateral striatal lesioning. These two factors may have reduced the impact of NA depletion and should be further investigated in future studies.

Our results show that DRN neurons are affected by depletion of both DA and NA, thus raising the possibility that non-motor symptoms in PD are a result of the intricate organization of DA and NA neuromodulation as well as the interactions between the different DRN neuronal populations. Moreover, our results highlight the complex interplay in the DRN between NA, DA, and 5-HT, but the precise pathophysiological processes resulting from loss of NA, and specifically the impact on DRN, are yet to be elucidated.

In conclusion, our study provides a quantitative description and classification scheme for two major neuronal populations in the DRN, DRN^5-HT^ and DRN^DA^ neurons. We identified novel electrophysiological and morphological changes in these populations in response to DA and NA depletion in the basal ganglia. Considering the involvement of DRN and LC in the development of non-motor comorbidities, this study provides useful insights to understand better how these areas are affected in the parkinsonian condition. Moreover, our data pave the way for future experiments to characterize these subpopulations in terms of receptor expression and synaptic connectivity to shed light on their functional roles particularly with regard to the wide variety of non-motor symptoms observed in PD.

## Methods

### Experimental model details

All animal procedures were performed in accordance with the national guidelines and approved by the local ethics committee of Stockholm, Stockholms Norra djurförsöksetiska nämnd, under ethical permits to G. F. (N12148/17, 14673-22) and G. S. (N2020/2022). All mice (N=43) were group-housed under a 12 hr light / dark schedule and given ad libitum access to food and water. Wild-type mice (‘C57BL/6J’, #000664, the Jackson laboratory) and DAT-cre (Stock #006660 the Jackson laboratory) mice crossed with homozygous tdTomato reporter mice (‘Ai9’, stock #007909, the Jackson laboratory) were used.

### 6-OHDA model

Three-months old, male and female C57BL/6J or DAT-tdTomato were deeply anesthetized with Isoflurane and mounted on a stereotaxic frame (Stoelting Europe, Dublin, Ireland). To achieve a partial striatal lesion, each mouse received a bilateral injection of 1.25 μl of 6-hydroxydopamine hydrochloride (6-OHDA, Sigma-Aldrich, 4μg/μl) or vehicle (0.9 % NaCl + Ascorbic Acid 0.02%) in the dorsolateral striatum, according to the following coordinates: anteroposterior +0.6 mm, mediolateral ± 2.2, dorsoventral -3.2 from Bregma, as previously described [48, 71]. One group of mice (referred to as DMI + 6-OHDA) was pre-treated with one injection of desipramine hydrochloride (DMI, Sigma-Aldrich, 25mg/kg i.p.) 30 minutes before the 6-OHDA infusion in order to protect the noradrenergic system [50].

For the LC lesion, mice received a unilateral injection of 1 μl of 6-OHDA (Sigma-Aldrich, 4μg/μl) or vehicle (0.9 % NaCl + Ascorbic Acid 0.02%) according to the following coordinates: anteroposterior –5.4 mm, mediolateral –0.9, dorsoventral -3.8 from Bregma.

### Slice preparation and electrophysiology

Three weeks after the 6-OHDA/vehicle injection, mice were deeply anaesthetized with isoflurane and decapitated. The brain was quickly removed and immersed in ice-cold cutting solution containing 205 mM sucrose, 10 mM glucose, 25 mM NaHCO3, 2.5 mM KCl, 1.25 mM NaH2PO4, 0.5 mM CaCl2 and 7.5 mM MgCl2. In all experiments, the brain was divided into two parts: the striatum was dissected from the anterior section for Western Blot and the posterior part was used to prepare coronal brain slices (250 μm) with a Leica VT 1000S vibratome. Slices were incubated for 30-60 min at 34°C in a submerged chamber filled with artificial cerebrospinal fluid (ACSF) saturated with 95% oxygen and 5% carbon dioxide. ACSF was composed of 125 mM NaCl, 25 mM glucose, 25 mM NaHCO3, 2.5 mM KCl, 2 mM CaCl2, 1.25 mM NaH2PO4, and 1 mM MgCl2. Subsequently, slices were kept for at least 60 min at room temperature before recording. Whole-cell patch clamp recordings were obtained in oxygenated ACSF at 35°C. Neurons were visualized using infrared differential interference contrast (IR-DIC) microscopy (Zeiss FS Axioskop, Oberkochen, Germany). DAT-tdTomato positive cells were identified by switching to epifluorescence using a mercury lamp (X-cite, 120Q, Lumen Dynamics). Up to three cells were patched simultaneously. Borosilicate glass pipettes (Hilgenberg) of 6 - 8 MOhm resistance were pulled with a Flaming / Brown micropipette puller P-1000 (Sutter Instruments). The intracellular solution contained 130 mM K-gluconate, 5 mM KCl, 10 mM HEPES, 4 mM Mg-ATP, 0.3 mM GTP, 10 mM Na2-phosphocreatine (pH 7.25, osmolarity 285 mOsm), 0.2% neurobiotin (Vector laboratories, CA) and Alexa-488 (75 μM) was added to the intracellular solution (Invitrogen). Recordings were made in current-clamp mode and the intrinsic properties of the neurons were determined by a series of hyperpolarizing and depolarizing current steps and ramps, enabling the extraction of sub- and suprathreshold properties. Recordings were amplified using MultiClamp 700B amplifiers (Molecular Devices, CA, USA), filtered at 2kHz, digitized at 10-20kHz using ITC-18 (HEKA Elektronik, Instrutech, NY, USA), and acquired using custom-made routines running on IgorPro (Wavemetrics, OR, USA). Throughout all recordings pipette capacitance and access resistance were compensated for and data were discarded when access resistance increased beyond 30 MOhm. Liquid junction potential was not corrected for.

### Quantification of electrophysiological parameters

Immediately after obtaining a whole-cell patch in DRN neurons, we first obtained a 10s voltage recording of the neural activity without injecting any current. This recording was used to calculate the average resting membrane potential in silent neurons and the firing frequency of spontaneously active neurons. Subsequently, neurons were held at –60mV while an extensive series of de- and hyperpolarizing current steps was applied. The amplitude of all current steps was scaled according to a test pulse that was set to evoke one to two AP. The resulting voltage recordings were used to extract and calculate the following parameters: The rheobase was defined as the minimum current required to evoke AP firing. AP parameters were extracted from recordings where DRN^5-HT^ neurons fired at 1 ± 0.3 Hz and DRN^DA^ neurons at 2 ± 0.3 Hz (i.e. close to their average spontaneous firing frequency) and values from individual APs were averaged. AP onset was extracted by quantifying where the rising slope of the AP (its first derivative) reached 5V/s and the end of the AP was defined as the time where the AP had repolarized to the same membrane voltage as found at the onset. The AP duration was calculated as the time between the onset and the offset. The amplitude of the AP was defined as the voltage difference between the onset and its peak. The amplitude of the AHP was defined as the voltage difference between the end of the AP and the subsequent local minimum. The end of the AHP was found by using a sliding window of 50ms to assess when the slope of the decaying AHP had first decreased to 0.005V/s or less. The AP drop rate was measured by injecting a current ramp into the neurons that evoked multiple APs. The amplitude of these APs was extracted as described above. The amplitude was plotted versus the injected current and a linear fit was applied whose slope constitutes the AP drop rate. The delay to the first spike constitutes the time between the onset of the current injection and the onset of the first AP in recordings. The input resistance was based on the slope of a linear fit across all current–voltage steps that resulted in a steady state voltage between –90 and –50mV (R = U / I). The steady state voltage was based on the average voltage found during a time window starting 0.5s after the beginning of a 1s long current step and lasting until the end of the current step. The amplitude of sag currents was defined as the average voltage difference between the steady state voltage and the peak voltage evoked by current steps that hyperpolarized the neurons to –90 ± 5mV. The peak voltage constituted the minimum voltage observed during the first 0.5s of the step. The time constant tau was extracted following injection of a 5ms long hyperpolarizing current step. We applied an exponential fit to the resulting voltage recording that started 1ms after the negative voltage peak had been reached and ended when the membrane potential had returned to the average baseline voltage preceding the step. Tau corresponds to K2 given the exponential fit is defined as y = K0 + K1*exp(-K2*x). Based on tau and the steady-state input resistance, we calculated the capacitance C according to C = tau / resistance. The IF slope was extracted from the linear fit applied to a current-frequency plot.

### Immunofluorescence

Following the recordings, slices were fixated overnight at 4°C in a 4% paraformaldehyde solution. Slices were then washed with PBS 1X. For the immunofluorescence, slices were treated with PBS 1X + Triton 0.3% and then incubated with a blocking solution of Normal Serum 10% and Bovine Serum Albumin 1% for 1h at room temperature. Afterward, slices were incubated overnight at 4°C with the following primary antibodies: rabbit anti-TH (Millipore, 1:1000), mouse anti-TPH (Sigma Aldrich, 1:600) and Streptavidin (Jackson Immunoresearch, 1:500). The following day, primary antibodies were washed out and slices were incubated with the appropriate fluorochrome-conjugated secondary antibodies.

For the immunostainings in the striatum, SNc, LC and cell counting in DRN, mice were deeply anesthetized and transcardially perfused with PFA 4%. The brains were extracted and post-fixed in PFA 4% for 24 hours. 40 μm coronal slices were prepared with a vibratome (Leica VT1000 S) and processed as described above.

### Confocal Microscopy Analysis

The slices were imaged using Confocal (ZEISS LSM 800) at 10X and 40X and z-stacks were retrieved. For cell identification, colocalization between neurobiotin and TH or TPH was evaluated.

### Morphological analysis

For morphological analysis of dendrites, the confocal z-stacks were used in a semi-manual reconstruction using neuTube [72] and custom code, as previously described [73]. Soma morphology was analyzed by tracing manually the cell body profile, excluding dendritic trunks, in order to measure area (μm^2^), perimeter, major and minor axis length (μm) and circularity values. Circularity, calculated as the ratio between the squared perimeter and the area (i.e. perimeter^2^/4π area), can be a value between 0 and 1 (1 for circular shapes and values < 1 for more complex shapes).

The morphological analysis was performed on the neurobiotin stacks.

### Western Blot

The striata were sonicated in 1% sodium dodecyl sulfate and boiled for 10 min. Equal amounts of protein (25 μg) for each sample were loaded onto 10% polyacrylamide gels and separated by electrophoresis and transferred overnight to nitrocellulose membranes (Thermo Fisher, Stockholm, Sweden). The membranes were immunoblotted with primary antibodies against actin (1:30,000, Sigma Aldrich, Stockholm, Sweden) and TH (1:2000, Millipore, Darmstadt, Germany). Detection was based on fluorescent secondary antibody binding (IR Dye 800CW and 680RD, Li-Cor, Lincoln, NE, USA) and quantified using a Li-Cor Odyssey infrared fluorescent detection system (Li-Cor, Lincoln, NE, USA). The TH protein levels were normalized for the corresponding actin detected in the sample and then expressed as a percentage of the control (Sham-lesion).

### Enzyme-linked immunosorbent assay

NA and 5-HT levels in the striatum were determined by Enzyme-linked immunosorbent assay (ELISA). Three weeks after the 6-OHDA injection, mice were killed by decapitation and the striatum was dissected out freehand on an ice-cold surface and weighted. The tissue was sonicated in a buffer with HCl 0.01M, EDTA 1mM and sodium metabisulfite 4 mM (25 μl/mg of tissue). The brain homogenates were centrifuged at 4 °C, 13000 rpm for 20 minutes and the supernatants were collected. The samples were assessed in analytic duplicate using Noradrenaline and Serotonin Research ELISA kits (LDN, Germany), according to the manufacturer’s instructions. The absorbance at 450 nm was measured using a microplate reader. Tissue concentrations of NA and 5-HT were determined using a standard curve.

### Statistical Analysis

Statistical analysis was performed using GraphPad Prism 9.2.0. Data were first tested for normality by Kolmogorov-Smirnov test. Two-groups analysis was performed by unpaired t-test for normally distributed data and the Mann-Whitney U test for non-normally distributed data. Three-groups analysis was performed by one-way ANOVA for normally distributed data or Kruskal-Wallis test for non-normally distributed data. Data are reported as average ± SEM. N indicates the number of mice, while n indicates the number of cells. Significance was set at p<0.05.

## Acknowledgments

We thank Elin Dahlberg for technical assistance and Kristoffer Tenebro Berglund for taking care of the mice. We also thank the members of the Silberberg and Fisone labs and the AND-PD consortium members for comments and discussions.

## Author contributions

Conceptualization: LB, YJ, RT, RM, GF, GS.

Investigation: YJ (electrophysiology, clustering analysis), LB (6-OHDA lesions, morphological analysis, cell counting, western blot, ELISA).

Visualization: YJ (electrophysiological and clustering results), LB (6-OHDA model, morphological reconstructions).

Supervision: GS and GF.

Writing—original draft: YJ and LB.

Writing—review & editing: LB, YJ, GF, GS.

## Data availability

All raw data is available upon reasonable request.

