## Supplementary Figures for "Serotonergic and dopaminergic neurons in the dorsal raphe are differentially altered in a mouse model for parkinsonism"

Supplementary Information

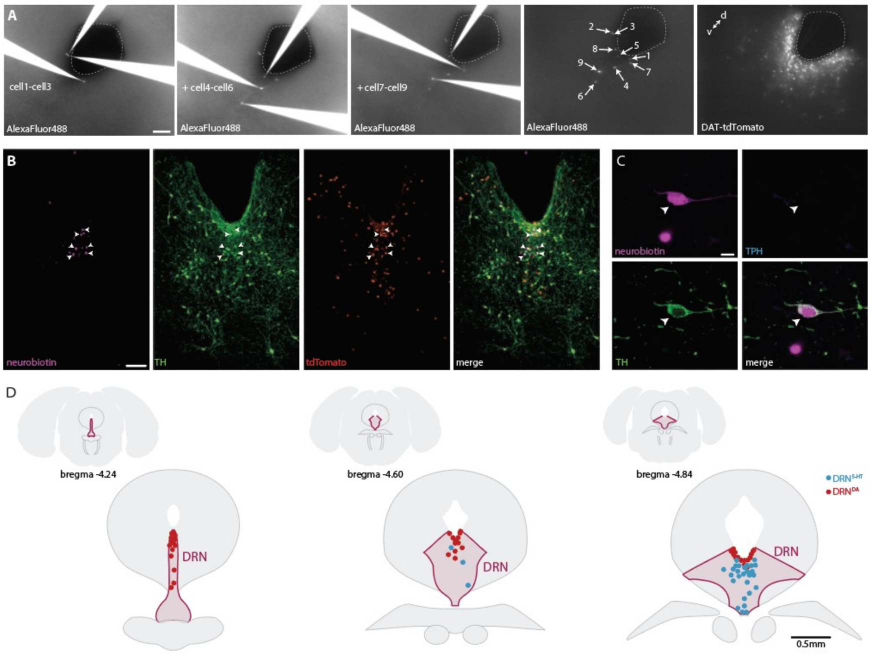


**Suppl. Fig. 1. Filling of neurons with AlexaFluor488 and neurobiotin for subsequent immunohistochemistry and topographical registration. (A)** Snapshots of recorded neurons filled with AlexaFluor488 facilitate post-hoc mapping of electrophysiology, anatomical location, and immunohistochemistry. Please note that cell 8 is shown at higher magnification in Fig. 1D. **(B)** Representative confocal pictures of post recording immunostaining for TH and neurobiotin revealing six TH-+ neurons (arrows) in a DAT-tdTomato mouse. **(C)** Post-recording immunostaining of a TH+ neuron in a wild-type mouse. **(D)** Histological verification of recording position. Representative schemes showing the location of $DRN^{5\text{-}HT}$ and $DRN^{DA}$ neurons recorded in Sham. Neurons are plotted in the nearest coronal section (-4.24, -4.60 or -4.84 from Bregma). Scale bars: A, B, 100 µm; C, 10 µm; D, 0.5 mm.

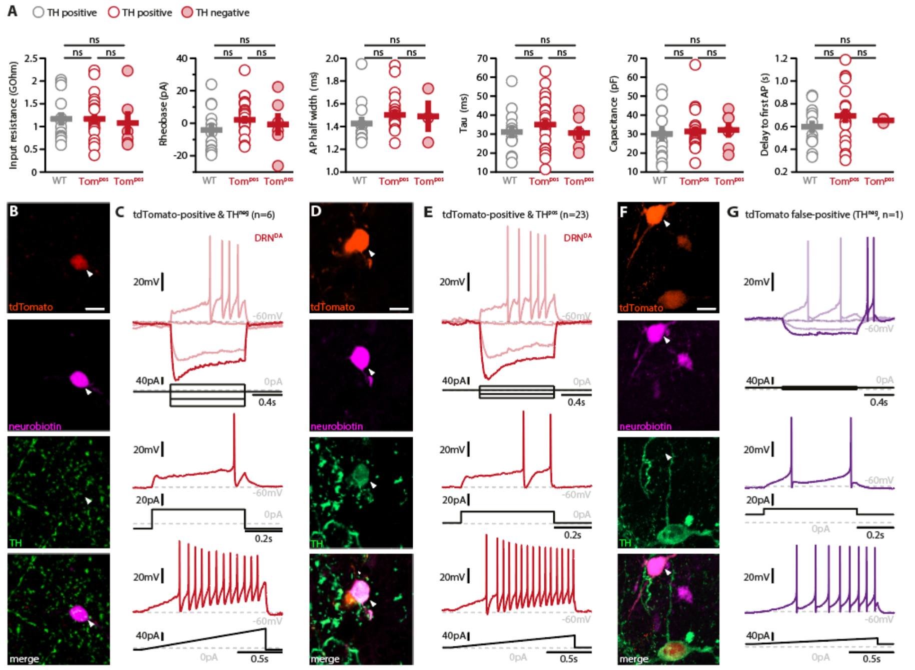


**Suppl. Fig. 2. Electrophysiological properties of DRN$^{DA}$ neurons recorded in wild-type or DAT-tdTomato mice do not differ. (A)** Intrinsic properties of DRN$^{DA}$ recorded in wild type (WT) or DAT-tdTomato (Tom$^{pos}$) mice (n = 13 DRN$^{DA}$ neurons from N = 5 wild type mice vs. n = 29 DRN$^{DA}$ neurons from N = 3 tdTomato mice). 6 of 29 tdTomato-+ neurons were found to be TH- (closed circles) while all other neurons were TH+ (open circles). No statistically significant differences were found when comparing the electrophysiological properties of these three groups (One-way Anova or Kruskal-Wallis test). **(B)** Post-recording immunohistochemistry showing the tdTomato-labelling, neurobiotin-filling and absence of TH staining in a DRN neuron (indicated with a white arrow). **(C)** Whole-cell recording of the neuron shown in (B) and its responses to current injections. Hierarchical cluster analysis of the electrophysiological profile suggests that this neuron is a DRN$^{DA}$ neuron despite the absence of TH staining. **(D)** Post-recording immunohistochemistry showing a DRN$^{DA}$ neuron whose identity is confirmed by both TH staining and tdTomato-labelling (indicated with a white arrow). **(E)** Whole-cell recording of the neuron shown in (D) showing the classic electrophysiological profile of DRN$^{DA}$ neurons. **(F)** Post-recording immunohistochemistry showing the tdTomato-labelling, neurobiotin-filling and absence of TH staining in a neuron false-positive for tdTomato (indicated with a white arrow). **(G)** Whole-cell recording of the neuron shown in (F) and its responses to different current injections. Please note the high input resistance, rebound spiking, biphasic afterhyperpolarisation and absence of spike amplitude accommodation. Scale bar: 10 µm.

**Deleted:** Mann Whitney U test

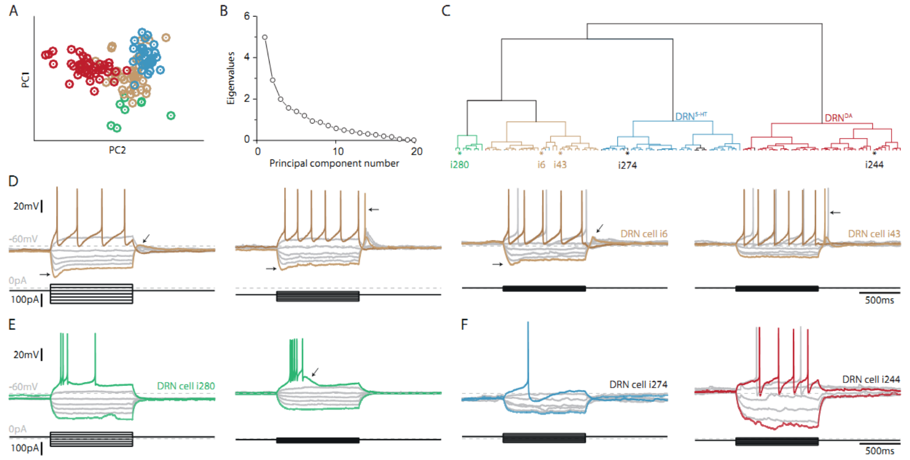


**Suppl. Fig. 3. Clustering of DRN neurons based on electrophysiological parameters. (A)** Principal component analysis (PCA) of 120 DRN neurons (N = 9) in Sham condition using 20 standard electrophysiological parameters. **(B)** Scree plot of the PCA. **(C)** Hierarchical cluster analysis (Ward's method, Euclidean distance) based on the first three principal components. Colors indicate four main clusters. TPH-+ ($DRN^{5\text{-}HT}$) neurons are indicated in blue, TH- and/or DAT-tdTomato positive ($DRN^{DA}$) neurons are indicated in red. Stars and corresponding identification numbers indicate DRN neurons of which example recordings are shown below. **(D-F)** Voltage responses to a series of current steps obtained from a variety of neurons in the DRN of Sham mice. **(D)** Four representative examples of the most frequently observed TH- and TPH- neurons recorded in the DRN. Please note the regular firing pattern, activation of sag currents (below -80mV), and rebound depolarization / spiking. **(E)** Two examples of DRN neurons characterized by plateau potentials, bursting and spike frequency accommodation. **(F)** Examples of molecularly unidentified neurons that were assigned to the $DRN^{5\text{-}HT}$ (left) and $DRN^{DA}$ (right) cluster by the hierarchical cluster analysis.

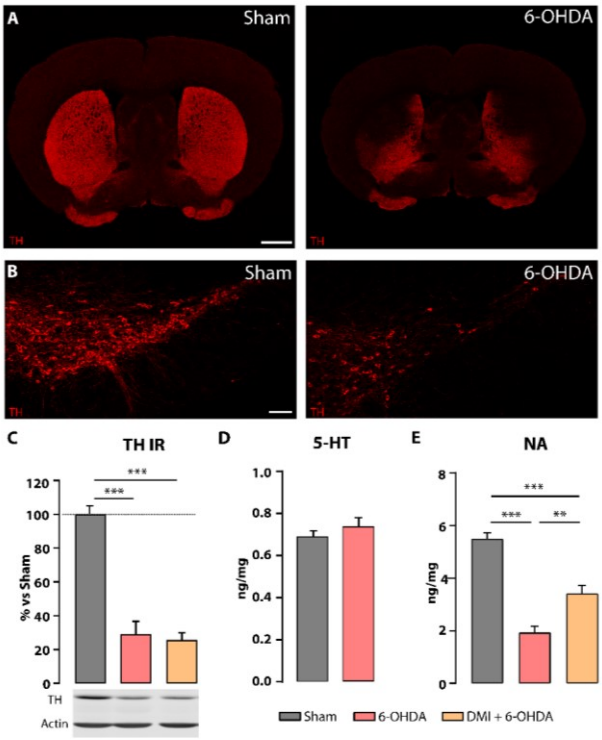


**Suppl. Fig. 4. The striatal injection of 6-OHDA induced 60-70 % TH loss in the striatum, did not alter striatal 5-HT levels but reduced striatal NA levels. The reduction of striatal NA levels was partially prevented by the pre-treatment with DMI. (A)** Representative confocal images in Sham (left) and 6-OHDA (right) showing the localization of TH loss in the dorsal striatum. **(B)** Representative confocal images in Sham (left) and 6-OHDA (right) showing the TH immunoreactivity in the SNc of Sham and 6-OHDA injected mice. **(C)** Top: bar chart showing the TH levels in Sham, 6-OHDA and DMI + 6-OHDA measured by Western Blot in the striatum (N=6-9 per group: one-way ANOVA). Data are normalized to Sham group (N=6-9 per group; one-way ANOVA). Bottom: representative blots of TH and Actin immunoreactivity in Sham (left), 6-OHDA (center) and DMI + 6-OHDA (right). **(D)** Bar chart showing the 5-HT levels in Sham and 6-OHDA measured by ELISA in the striatum. (N=5-6 per group). **(E)** Bar chart showing the NA levels in Sham, 6-OHDA and DMI + 6-OHDA measured by ELISA in the striatum (N=9-10 per group; one-way ANOVA). Data are presented as mean ± SEM. ***p<0.001, **p<0.01. Scale bars: A, 1 mm; B, 100 µm.

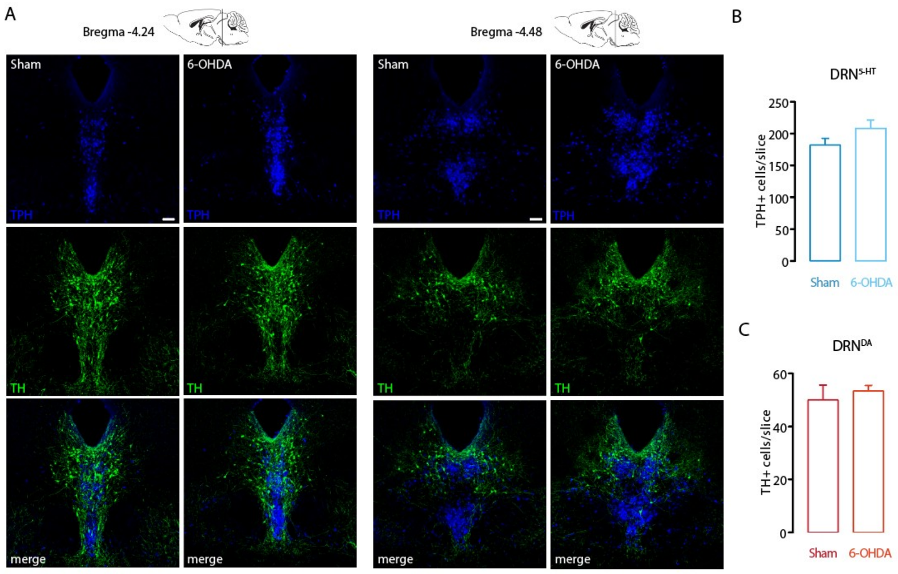


**Suppl. Fig. 5. The 6-OHDA injection did not affect the number of DRN$^{5\text{-}HT}$ and DRN$^{DA}$ neurons. (A)** Representative confocal pictures of DRN in Sham and 6-OHDA mice at different antero-posteriorities. **(B)** Bar chart showing the density of TPH+ neurons in the DRN in Sham and 6-OHDA groups. **(C)** Bar chart showing the density of TH+ neurons in the DRN in Sham and 6-OHDA groups. N=4 per group. Data are shown as mean ± SEM. Scale bar 100 µm.

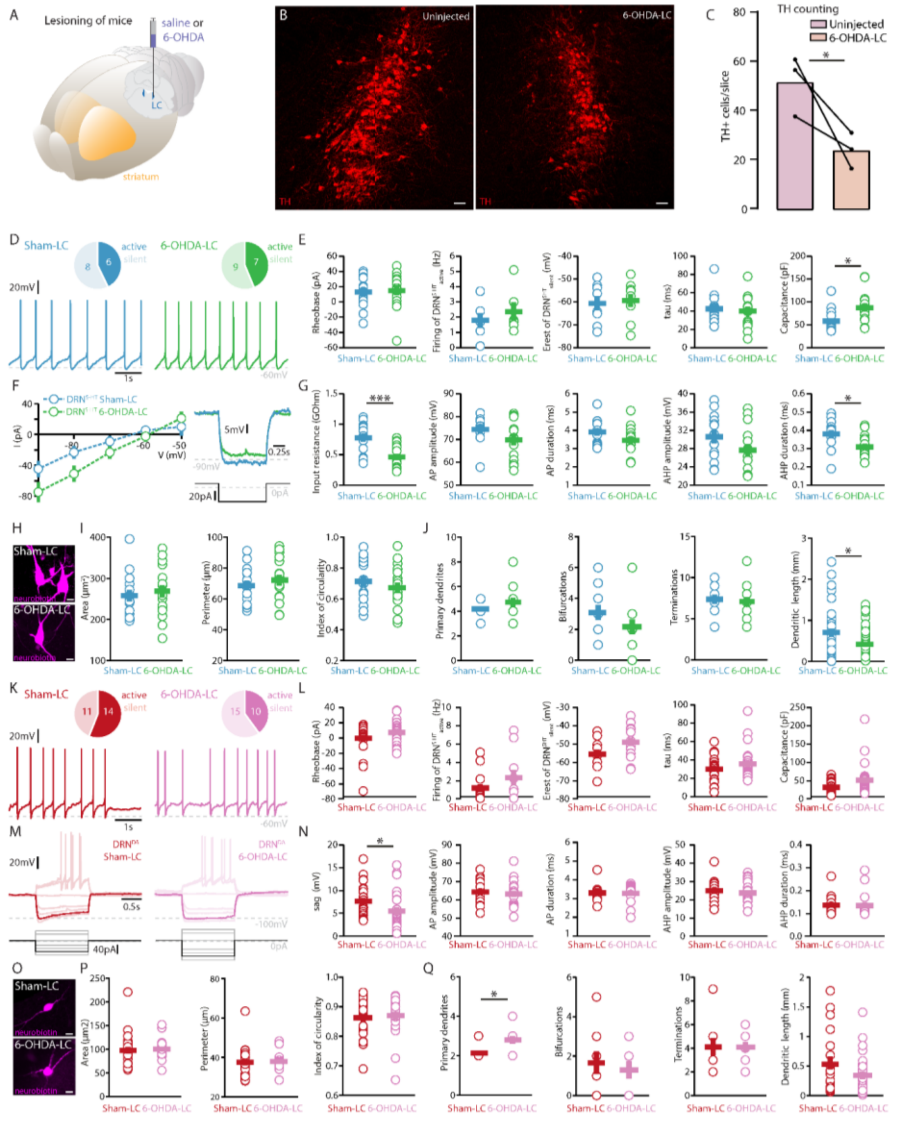


**Suppl. Fig. 6. Selective lesioning of the NA system based on 6-OHDA injections in the LC affects DRN neurons mildly. (A)** Scheme showing unilateral injection of vehicle / 6-OHDA in LC. **(B)** Representative confocal images of the lesion induced by the unilateral injection of 6-OHDA in the LC (left: uninjected side, contralateral to the 6-OHDA injection; right: 6-OHDA-injected side). **(C)** Bar chart showing the density of TH+ neurons in the LC in the uninjected and 6-OHDA-injected sides. Data are shown as mean

**Formatted:** English (United Kingdom)

± SEM (N=3 per group, unpaired t-test). **(D)** Top: pie charts showing the number of spontaneously active (dark) and silent (pale) $DRN^{5-HT}$ neurons in mice injected with vehicle (Sham-LC, left) and mice injected with 6-OHDA in LC (6-OHDA-LC, right). Bottom: representative recordings of spontaneously active $DRN^{5-HT}$ neurons (I = 0 pA). **(E)** Quantification of the rheobase (Sham-LC: n = 14, 6-OHDA-LC: n = 16), the firing frequency of spontaneously active cells (Sham-LC: n = 6, 6-OHDA-LC: n = 7), the resting membrane potential of silent $DRN^{5-HT}$ neurons (Sham-LC: n = 8, 6-OHDA-LC: n = 9), the time constant tau and the capacitance (Sham-LC: n = 14, 6-OHDA-LC: n = 15) **(F)** Representative responses of $DRN^{5-HT}$ neurons to hyper- (left) and depolarizing (right) current steps. **(G)** Quantification of the input resistance (Sham-LC: n = 14, 6-OHDA-LC: n = 16) and AP properties of $DRN^{5-HT}$ neurons (AP and AHP amplitude and duration: Sham-LC: n = 13, 6-OHDA-LC: n = 15). **(H)** Representative confocal pictures of soma from $DRN^{5-HT}$ neurons in Sham-LC (top) and 6-OHDA-LC mice (bottom) **(I)** Morphological descriptors of the soma size and shape in $DRN^{5-HT}$ neurons (Sham-LC: n = 15, N = 5, 6-OHDA-LC: n = 17, N = 5) **(J)** Morphological descriptors of the dendritic tree in $DRN^{5-HT}$ neurons. (Sham-LC: n = 11, N = 4; 6-OHDA-LC: n = 12, N = 4; Mann-Whitney U test) **(K)** Same as in (D) for $DRN^{DA}$ neurons. **(L)** Quantification of the rheobase (Sham-LC: n = 25, 6-OHDA-LC: n = 25), the firing frequency of spontaneously active cells (Sham-LC: n = 14, 6-OHDA-LC: n = 10), the resting membrane potential of silent $DRN^{DA}$ neurons (Sham-LC: n = 11, 6-OHDA-LC: n = 15), the time constant tau and the capacitance (Sham-LC: n = 25, 6-OHDA-LC: n = 25). **(M)** Representative responses of $DRN^{DA}$ neurons recorded in Sham-LC (left) and 6-OHDA-LC (right) to current steps. The step hyperpolarizing the neurons to –100mV is highlighted. **(N)** Quantification of the sag amplitude (Sham-LC: n = 23, 6-OHDA-LC: n = 23) and AP properties of $DRN^{DA}$ neurons (AP and AHP amplitude and duration: Sham-LC: n = 15, 6-OHDA-LC: n = 18). **(O)** Representative confocal pictures of soma from $DRN^{DA}$ neurons in Sham-LC (top) and 6-OHDA-LC mice (bottom) **(P)** Morphological descriptors of the soma size and shape in $DRN^{DA}$ neurons (Sham-LC: n = 23, N = 5, 6-OHDA-LC: n = 23, N = 5) **(Q)** Morphological descriptors of the dendritic tree in $DRN^{DA}$ neurons (Sham-LC: n = 10, N = 3; 6-OHDA-LC: n = 10, N = 4; Mann-Whitney U test). All electrophysiological data was acquired in N = 5 Sham-LC and N = 5 6-OHDA-LC mice and statistics are based on unpaired t-test or Mann Whitney U test). Data are shown as mean ± SEM, * $p < 0.05$, ** $p < 0.01$, *** $p < 0.001$.

Scale bars: B, 50 µm, H-O, 10 µm
